# Tissue culture as a source of replicates in non-model plants: variation in cold tolerance in *Arabidopsis lyrata* ssp. *petraea*

**DOI:** 10.1101/043695

**Authors:** Tanaka Kenta, Jessica E.M. Edwards, Roger K. Butlin, Terry Burke, W. Paul Quick, Peter Urwin, Matthew P. Davey

## Abstract

Whilst genotype–environment interaction is increasingly receiving attention by ecologists and evolutionary biologists, such studies need genetically homogeneous replicates—a challenging hurdle in outcrossing plants. This could potentially be overcome by using tissue culture techniques. However, plants regenerated from tissue culture may show aberrant phenotypes and “somaclonal” variation. Here we examined the somaclonal variation due to tissue culturing using the response of the photosynthetic efficiency (chlorophyll fluorescence measurements for *F*_*v*_/*F*_*m*_, *F*_*v*_’/*F*_*m*_’ and Φ_PSII_, representing maximum efficiency of photosynthesis for dark‐ and light-adapted leaves, and the actual electron transport operating efficiency, respectively) to cold treatment, compared to variation among half-sibling seedlings from three different families of *Arabidopsis lyrata* ssp. *petraea*. Somaclonal variation was limited and we could successfully detect within-family variation in change in chlorophyll fluorescence by cold shock with the help of tissue-culture derived replicates. Icelandic and Norwegian families exhibited higher chlorophyll fluorescence, suggesting higher cold tolerance, than a Swedish family. Although the main effect of tissue culture on *F*_*v*_/*F*_*m*_, *F*_*v*_’/*F*_*m*_’ and Φ_PSII_ was small, there were significant interactions between tissue culture and family, suggesting that the effect of tissue culture is genotype–specific. Tissue-cultured plantlets were less affected by cold treatment than seedlings, but to a different extent in each family. These interactive effects, however, were comparable to, or much smaller than the single effect of family. These results suggest that tissue culture is a useful method for obtaining genetically homogenous replicates for studying genotype–environment interaction related to adaptively relevant phenotypes, such as cold tolerance, in non-model outcrossing plants.

## Introduction

Genotype–environment interaction on a phenotype or reaction norm may modulate natural selection (Wright 1931; Sultan 1987). The genetic basis of genotype–environment interaction is increasingly receiving attention (El-Soda *et al*. 2014; Yap *et al*. 2011); however, such advances have been concentrated in inbreeding organisms such as *Arabidopsis thaliana* (e.g. Bloomer *et al*. 2014; El-Soda *et al*. 2014; Sasaki *et al*. 2015; Stratton 1998) and *Caenorhabditis elegans* (Gutteling *et al*. 2007), because genetically isogenic individuals permit a given genotype to be exactly repeated in multiple environments. Recently, the wild relatives of model organisms are increasingly being exploited by evolutionary biologists to understand adaptation and speciation (Clauss & Koch 2006; Mitchell-Olds 2001). However, one disadvantage of non-model plants with outcrossing mating systems is that they cannot usually be exploited to produce the genetically homogeneous or inbred recombinant lines that enable researchers to study the reaction norms of a single genotype in multiple environments (Dorn *et al*. 2000) or to map novel QTLs in previously-genotyped lines (Alonso-Blanco *et al*. 2005). This disadvantage could be compensated for by using cutting techniques to produce multiple clones from single genotypes (Sultan & Bazzaz 1993; Waitt & Levin 1993; Wu 1998). This method is only applicable to plants capable of vegetative propagation, and it also needs relatively large plant bodies to produce many replicate clones. Another technique applicable to a wider range of plants with relatively small starting plant material is tissue culture (George & Sherrington 1984). However, tissue culture has been exploited rarely for studies on the genetic basis of genotype–environment interaction, and the few existing studies (Glock 1989; Glock & Gregorius 1986) focused only on callus characteristics as target phenotypes. One potential issue that should be carefully considered is that tissue-culture derived microshoots can express phenotypic, “somaclonal” variation (Larkin & Scowcroft 1981) or may sometimes show aberrant morphology and physiology *in vitro* (Joyce *et al*. 2003). This somaclonal variation resembles that induced by physical mutagens, with elevated levels of chromosome breakage and rearrangement, polyploidy, aneuploidy, transposon activation and point mutation (D'Amato & Bayliss 1985). Therefore, with a view to exploiting the techniques of tissue culturing more widely in studies of genotype–environment interaction in outcrossing plants, it is necessary to extend our knowledge on how propagation by tissue culture generates variation in phenotypes that are relevant to adaptation in natural environments, compared to other sources of genetically-related replicates such as outbred siblings.

Key plant properties that have attracted marked attention in the field of adaptation to various environments are stress tolerances (e.g. Hong & Vierling 2000; Kwon *et al*. 2007; Lexer *et al*. 2003; Quesada *et al*. 2002; Steponkus *et al*. 1998; Zhang *et al*. 2004; Zhen & Ungerer 2008). One trait that can be used to indicate tolerance against various physical stressors in plants is photosynthetic performance. Photosystem II (PSII) activity is sensitive to both biotic and physical environmental factors (Murchie & Lawson 2013). Chlorophyll fluorescence can be used to determine the maximum efficiency with which light absorbed by pigments of photosystem II (PSII) is used to drive photochemistry in dark‐ (*F*_*v*_/*F*_*m*_) or light‐ (*F*_*v*_’/*F*_*m*_) adapted material and the operating efficiency of PSII (Φ_PSII_). It is a reliable indicator of photoinhibition and damage to the photosynthetic electron transport system (Maxwell & Johnson 2000; Quick & Stitt 1989). Changes in chlorophyll fluorescence have been successfully used in *Arabidopsis thaliana* to quantify tolerance to cold and freezing temperatures (Ehlert & Hincha 2008; Heo *et al*. 2014; Mishra *et al*. 2014), drought (Bresson *et al*. 2015; McAusland *et al*. 2013; Woo *et al*. 2008), and salt and heavy-metal stress (Yuan *et al*. 2013), as well as in various other plants for tolerance to cold and freezing temperatures (Baldi *et al*. 2011; Khanal *et al*. 2015; Medeiros *et al*. 2012; Xie *et al*. 2015), drought (Jansen *et al*. 2009) and salt (Yuan *et al*. 2013). If variation in chlorophyll fluorescence can be properly estimated using tissue-culture derived clones, therefore, it would enhance studies in genotype–environment interaction for stress tolerance in outcrossing plants.

To this end, we have studied change in chlorophyll fluorescence following cold shock in a wild relative of a model plant species. *Arabidopsis lyrata* ssp. *petraea* is a close relative of the model species *A. thaliana,* but with a different ecology, life history and population genetics (Charlesworth *et al*. 2003; Davey *et al*. 2008; Davey *et al*. 2009; Kuittinen *et al*. 2008; Kunin *et al*. 2009). Whilst *A. thaliana* is mainly selfing, with a low level of genetic diversity within a population, *A. lyrata* ssp. *petraea* is outcrossing, with a high level of genetic diversity even within a population (Clauss & Mitchell-Olds 2006; Heidel *et al*. 2006; Kunin *et al*. 2009; Schierup *et al*. 2008). Further studies on genetic and phenotypic variation in spatially distinct individuals and in closely-related plants will clarify whether or not locally advantageous alleles are fixed and if local populations are in evolutionary equilibrium, and are thus important in our understanding of the evolutionary responses to environmental change. Distinguishing phenotypic variation among closely related individuals from measurement errors is difficult; however, this becomes possible if we can quantify the error within the same genotype using tissue-cultured clones.

In this study, we measured the chlorophyll fluorescence parameters *F*_*v*_/*F*_*m*_, *F*_*v*_’/*F*_*m*_’ and Φ_PSII_ before and after cold shock, as an index of cold tolerance, for seedlings from three families from geographically isolated populations of *A. lyrata* ssp. *petraea*, and tissue cultured plantlets derived from several genotypes (seeds) in each of those families (Table 1). In order to evaluate the usefulness of tissue culture for obtaining genetically homogenous replicates and to assess how much adaptively-relevant variation exists within the species, we tested whether (i) among-genotype phenotypic variation could be detected with the help of replication of tissue cultured plantlets, (ii) somaclonal variation would remain in the range of other components of variation such as within-family variation of seedlings, (iii) phenotypic variation in adaptively relevant traits would exist between families and (iv) tissue-culturing affected these measurements of chlorophyll fluorescence.

**Table 1.**
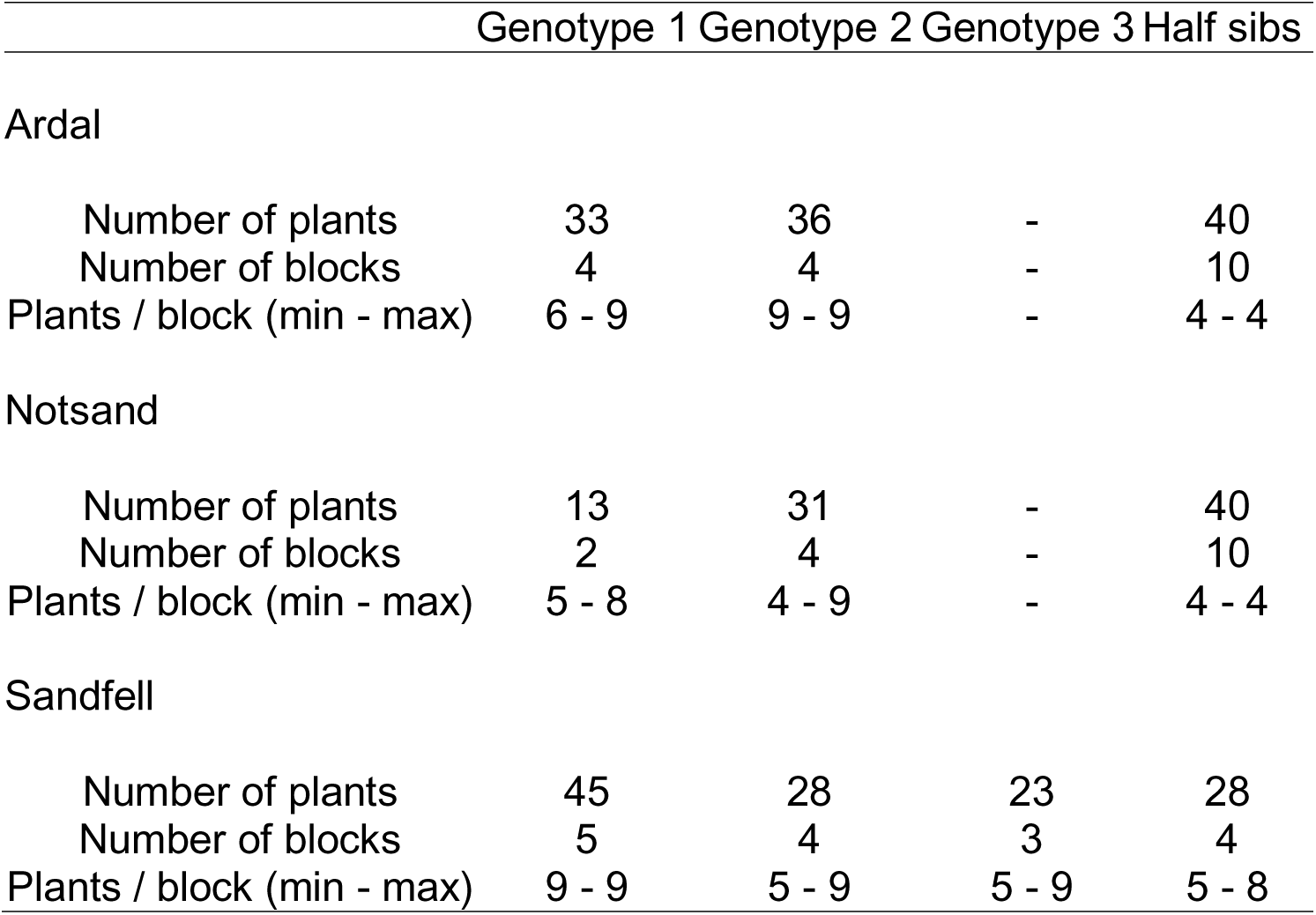
Numbers of plants and blocks in each family (Ardal, Notsand and Sandfell). Plants were either seedlings in a half-sibling family or tissue-cultured clonal plantlets from genotypes derived from a seed from each family.

## Material and Methods

### Plants

Seeds of *Arabidopsis l. petraea* were collected from geographically separated populations in Ardal (Norway) (61°19′25″N, 7°50′00″E, alt. 63 m), Notsand (Sweden) (62°36′31″N, 18°03′37″E, alt. 3 m) and Sandfell (Iceland) (64°04′14″N, 21°41′06″E, alt. 123 m). No specific permits were required for the seed collection for this study because these locations were not privately owned or protected in any way and because the species was not protected in these countries. The species is a perennial herb and keeps leaves throughout the year. We used a family of seeds that were at least half-siblings, from one mother plant in each population. We grew 28–40 seedlings per family and in each case derived 44–69 tissue-cultured plantlets from 2–3 seeds (1 genotype = cloned plantlets from one seed) of each family.

### Tissue culture

Seeds were sterilised in 10% commercial bleach for 20 min, washed in sterile water and stored at 4°C overnight. The seeds were then placed onto 50% strength Murashige and Skoog (MS) medium (Melford Laboratories Ltd, Ipswich, UK), pH 5.7, supplemented with 1 % sucrose, 5 mg/L silver thiosulphate and solidified with 1 % plant agar (Melford Labs. Ltd). The agar plates were held vertically, allowing for maximum recovery of root tissue. After 4 weeks the root systems were excised and placed intact onto Callus Induction Medium (CIM) (Clarke *et al*., 1992) solidified with 0.55% plant agar. Plates were incubated at 23 °C for 3 days then the roots were cut into 5 mm lengths and placed in bundles on fresh CIM plates that were further incubated at 20°C for 2–3 days. The root sections from each plant were resuspended in 10 ml molten Shoot Overlay Medium (SOM) (Clarke *et al*., 1992) solidified with 0.8 % low gelling-temperature agarose and poured over a single 90 mm plate of Shoot Induction Medium (SIM) (Clarke *et al*., 1992) solidified with 0.55 % plant agar and lacking antibiotics. The plates were incubated at 20 °C under a 16-hour day length. Once shoots started to form from the calli they were transferred to 50 % strength MS medium, pH5.7, supplemented with 1 % sucrose and solidified with 0.55% plant agar, such that each plate contained 9 clones of the same genotype. A total of 4–9 plantlets survived per plate. Each plate was treated as a block in the following experiment.

### Seedling growth

Seeds were sown in Levington M3 compost within individual plug trays. Families were randomised within each tray and trays were randomly repositioned every other day. Plants were watered from the base of the pot as required with reverse-osmosis (RO) purified water. No additional nutrients were added to the soil or water. Plants were established to 6–8 leaf stage in controlled-environment growth cabinets (Conviron Controlled Environments Limited, Canada) set to a 12/12 hour day/night cycle, 20/15 °C day/night, 70 % humidity; atmospheric CO_2_ concentration was 400 ppm and photosynthetically-active radiation 250 μmol m^-2^ s^-1^. Chlorophyll fluorescence measurements were taken just prior to and after a 24 hour cold treatment in which plants were exposed to the same conditions as above, apart from the temperature being decreased to 3 °C. 5–8 seedlings from the same family were treated as a block in the following experiment.

### Chlorophyll fluorescence

Pre-cold and post-cold treatment measurements of chlorophyll fluorescence were obtained using a chlorophyll fluorescence imager using Fluorimager software (Technologica Ltd., Colchester, UK). Each block of plants was dark adapted for at least 15 minutes before the maximum efficiency of photosystem II (*F*_*v*_/*F*_*m*_) was measured to a blue light pulse at 3000 μmol m^-2^ s^-1^ for 200 ms. Following this pulse, the plants were exposed to an actinic light of 150 μmol m^-2^ s^-1^ for six minutes, followed by pulses of 3000 μmol m^-2^ s^-1^ for 200 ms to obtain measures of maximum efficiency of photosystem II (*F*_*v*_’/*F*_*m*_) of light-adapted plant material and the operating efficiency of photosystem II (Φ_PSII_) in light-adapted plant material. Mean values of *F*_*v*_/*F*_*m*_, *F*_*v*_’/*F*_*m*_’ and Φ_PSII_ for each plant were taken from the image of each whole plant.

All these phenotypic data are available in Dryad Digital Repository: http://dx.doi.org/10.5061/dryad.xxxxx.

### Statistical analyses

To examine the relative importance of among-family and among-genotype variation in cold tolerance, we used nested ANOVA to partition the total variance in the difference in each chlorophyll fluorescence measurement (*F*_*v*_/*F*_*m*_, *Fv*’/*Fm*’ or Φ_PSII_) induced by cold shock:

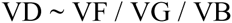

where VD was the total variance in difference in each type of chlorophyll fluorescence for a plant individual between two measurements (i.e. value after cold shock minus that before cold shock), VF was the component of among-family variance, VG was the component of among-genotype variance nested in VF and VB was the component of among-block variance nested in VG. We did this analysis separately for the tissue-cultured plants and seedlings, in order to evaluate variation in each natural and tissue-cultured condition. The VG term was not applied to the analysis for seedlings. We also conducted variance component analysis using the varcomp function in the ape library and the lme function using *R* 2.8.0(R Development Core Team 2008).

We tested whether variance in the change of *F*_*v*_/*F*_*m*_, *Fv*’/*Fm*’ or Φ_PSII_ due to cold shock among tissue-culture derived plantlets within each genotype was different from that in seedlings of half-siblings of the same family using Bartlett tests. Because the number of blocks differed between seedlings and tissue-cultured plantlets (Table 1), we checked first whether the difference in the number of blocks affected the variance, by re-sampling all possible combinations of 4 blocks from the 10 blocks of halfsiblings in Ardal and Notsand. Reducing block number changed the original variance for 10 blocks only < ±3 % without systematic bias.

Finally, we evaluated the effect of several factors on each type of chlorophyll fluorescence measurement before and after cold treatment. We constructed the following linear mixed-effect model, in which plant individual was treated as a random effect:

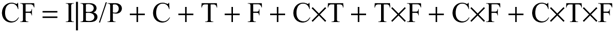

where CF was a single measurement of either *F*_*v*_/*F*_*m*_, *Fv*’/*Fm*’ or Φ_PSII_ and I|B/P was the intercept with random effects of block, and individual plant nested in each block, C was a binary variable of cold shock (1 for shocked and 0 for not), T was a binary variable of tissue culture (1 for tissue cultured and 0 for not) and F was a categorical variable of family (3 families), followed by the interaction terms among those variables. The effect of each term was estimated by the lme function using the statistical software *R* 2.8.0 (R Development Core Team 2008). Akaike's Information Criterion (AIC) was compared between the full model and a model lacking each term in a stepwise manner and the best model with the lowest AIC was selected, followed by testing the significance of each selected parameter using the Wald test.

## Results

### Variance components in cold-response of *Fv/Fm Fv’/Fm’* and Φ_PSII_

In the seedlings, the changes in *F*_*v*_/*F*_*m*_, *Fv*’/*Fm*’ or Φ_PSII_ by cold treatment varied significantly among families, explaining 4.9–9.1 % of the total variance (Table 2). For the tissue-cultured plantlets, the change in those indices by cold treatment did not vary significantly among families, but did vary significantly among genotypes within family, this component explaining 8.5–31.5 % of the total variance. The within-block variance component for tissue-cultured plantlets was 61.7–81.8 % and tended to be smaller than this component for seedlings (89.1–92.2 %).

### Evaluation of somaclonal variation in comparison to within-family variation

Variances in the change of *F*_*v*_/*F*_*m*_, *Fv*’/*Fm*’ or Φ_PSII_ among clones within genotype were clearly smaller than those among half-siblings of the same family in the Sandfell family. Most genotypes had significantly smaller variances in *F*_*v*_/*F*_*m*_, *F*_*v*_’/*F*_*m*_’ and Φ_PSII_ than half-sibs as shown by the Bartlett test (Fig. 1). Similar patterns were observed in Notsand and Ardal. No studied genotype had larger variance among clones than the variance among half-siblings in any family.

**Figure. 1.**
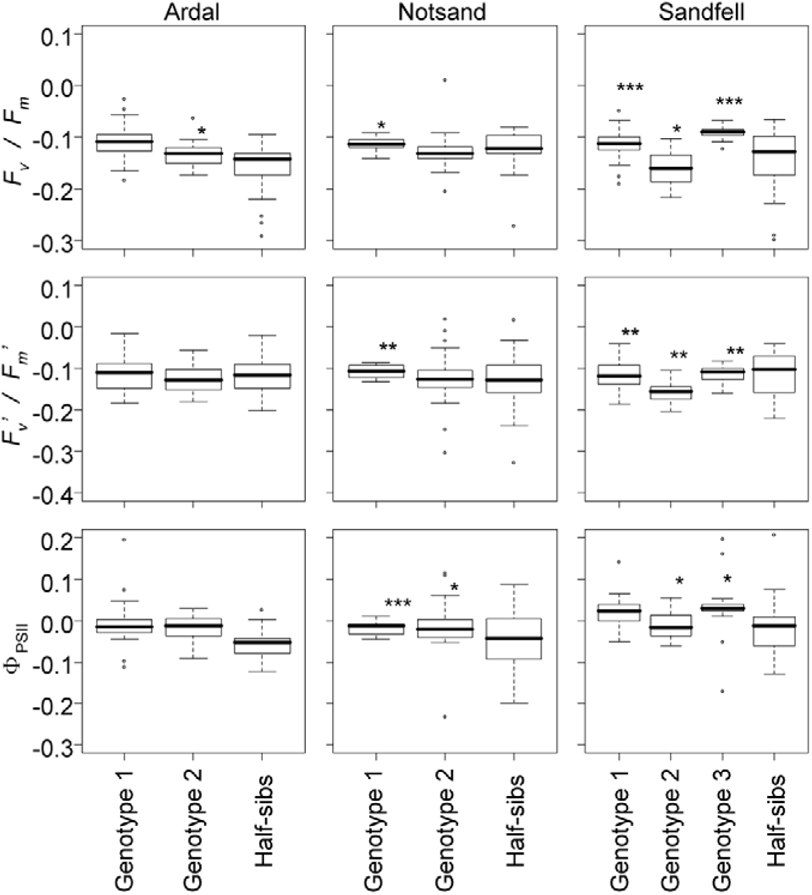
Change in chlorophyll fluorescence (*F*_*v*_/*F*_*m*_, *F*_*v*_’/*F*_*m*_’ and Φ_PSII_) in seedlings or plantlets originating from Norway (Ardel), Sweden (Notsand) and Iceland (Sandfell) after cold-treatment *(values after shock – those before shock)*. *, ** *and* *** = *P < 0.05, P* < 0.01 and *P* < 0.001, respectively, (Bartlett test) indicate a significantly lower variance of the genotype than among half-siblings in the same family. Three *F*_*v*_/*F*_*m*_ values (0.340, 0.375, 0.592) and an *F*_*v*_’/*F*_*m*_’ value (0.354) in Sandfell half-siblings were out of the vertical ranges shown but were included in the statistical tests.

### Effects of cold shock, tissue culturing and family on *Fv*/*Fm Fv’*/*Fm’*and Φ_PSII_

All single effects of cold shock, tissue culture and family and all possible interaction combinations among them affected *F*_*v*_/*F*_*m*_ and *F*_*v*_*’*/*F*_*m*_*’*, and all such effects except the 3-way interaction between cold shock, tissue culture and family affected Φ_PSII_, according to the best model (Table 3) based on Akaike's Information Criterion (AIC). Cold shock and family were the strongest single effects. The interaction between these two factors was also found to change all three measurements of chlorophyll fluorescence, indicating that the effect of cold shock depended on family. The effect of tissue culture was relatively small and not significant for any of the chlorophyll fluorescence measures. We found substantial interactions between tissue culture and family and interactions among cold shock, tissue culture and family, indicating that the effect of tissue culture depended on family.

## Discussion

### Among-genotype variance

We were able to test for among-genotype variance using replicates generated by tissue culture within genotypes and we detected such variance in *F*_*v*_/*F*_*m*_, *F*_*v*_*’*/*F*_*m*_*’* and Φ_PSII_ measurements (Table 2). On the other hand, we showed significant but low somaclonal variation. The within-block variance component for tissue-cultured plantlets was relatively small (Table 2). The Bartlett tests showed that somaclonal variation was smaller than, or at least remained within the range of, the within-family variance, which is the smallest naturally observed component of variation in the hierarchy of genetic structure (Fig. 1). In *A. thaliana*, studies of natural variation have focused mainly on between-population variation (e.g. (Shindo *et al*. 2007). In contrast, *A. lyrata* has substantial within-population variation, for example in the composition of glucosinolates (Clauss *et al*. 2006) or self-incompatibility genes (Schierup *et al*. 2008). In this paper, we showed that there is within-family as well as among-family, and thus among-population, genetic variation in *A. lyrata* ssp. *petraea*. Within-family genetic variance was relatively large in Sandfell (Iceland). The observed within-family genetic variances in adaptively relevant traits highlight the wide potential for evolutionary adaptation of the species and further validates the usefulness of relatives of model organisms in evolutionary biology (Clauss & Koch 2006; Mitchell-Olds 2001).

**Table 2.**
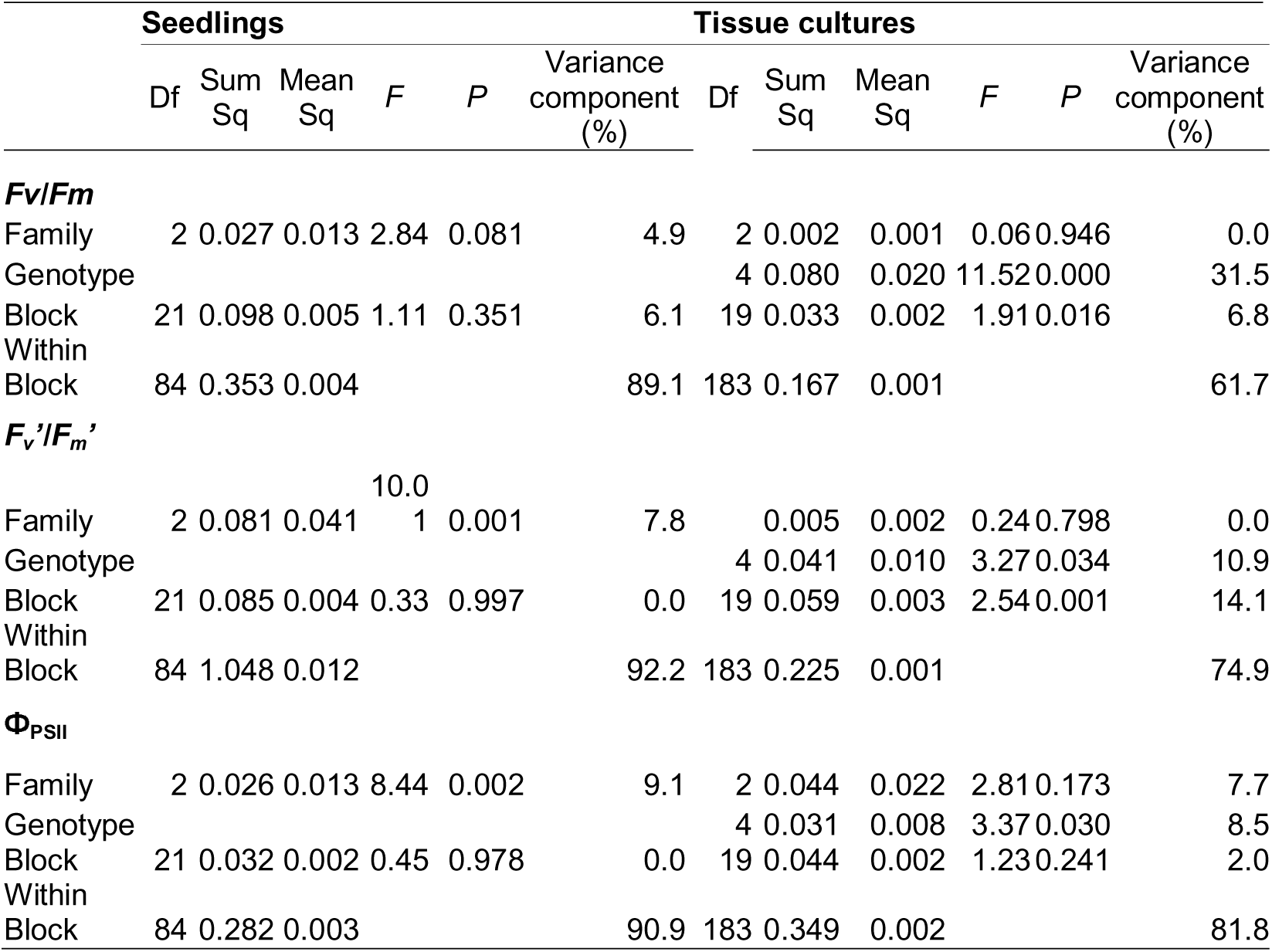
Analysis of variance for change in *F*_*v*_/*F*_*m*_, *F*_*v*_’/*F*_*m*_’ and Φ_PSII_ by cold treatment for non-tissue cultured seedlings and tissue-cultured plantlets. Family and Block refer to variation among families and among blocks within families, respectively

### Among-family variance

There was significant or marginally significant among-family variance in the change of *F*_*v*_/*F*_*m*_, *F*_*v*_*’*/*F*_*m*_*’* and Φ_PSII_ values by cold treatment for seedlings (Table 2). In *A. thaliana*, the change in chlorophyll fluorescence from before to after cold shock correlates with tolerance to sub-zero temperatures measured by electrolyte leakage or survival and, therefore, this is regarded as an indicator of cold tolerance (Ehlert & Hincha 2008; Heo ei *al*. 2014; Khanal *et al*. 2015). Therefore, our result also represents evidence for among-family (thus possibly among-population) variance in cold tolerance. Linear mixed models (Table 3) showed that *F*_*v*_*’*/*F*_*m*_*’* after cold-shock was higher in family Ardal and Sandfell, and *F*_*v*_/*F*_*m*_ and Φ_PSII_ after cold-shock was higher in family Sandfell, compared to Notsand (Sweden). These results are consistent with families Aradal (Norway) and Sundfell (Iceland) being derived from relatively high latitude and high altitude and so having high cold tolerance. This among-family effect was weaker for tissue-cultured plantlets (Table 3). This may be due to the small number of genotypes for each family in our nested experimental design, or, could be explained by the main part of the among-family variance detected for seedlings being due to among-genotype variance within families.

**Table 3.**
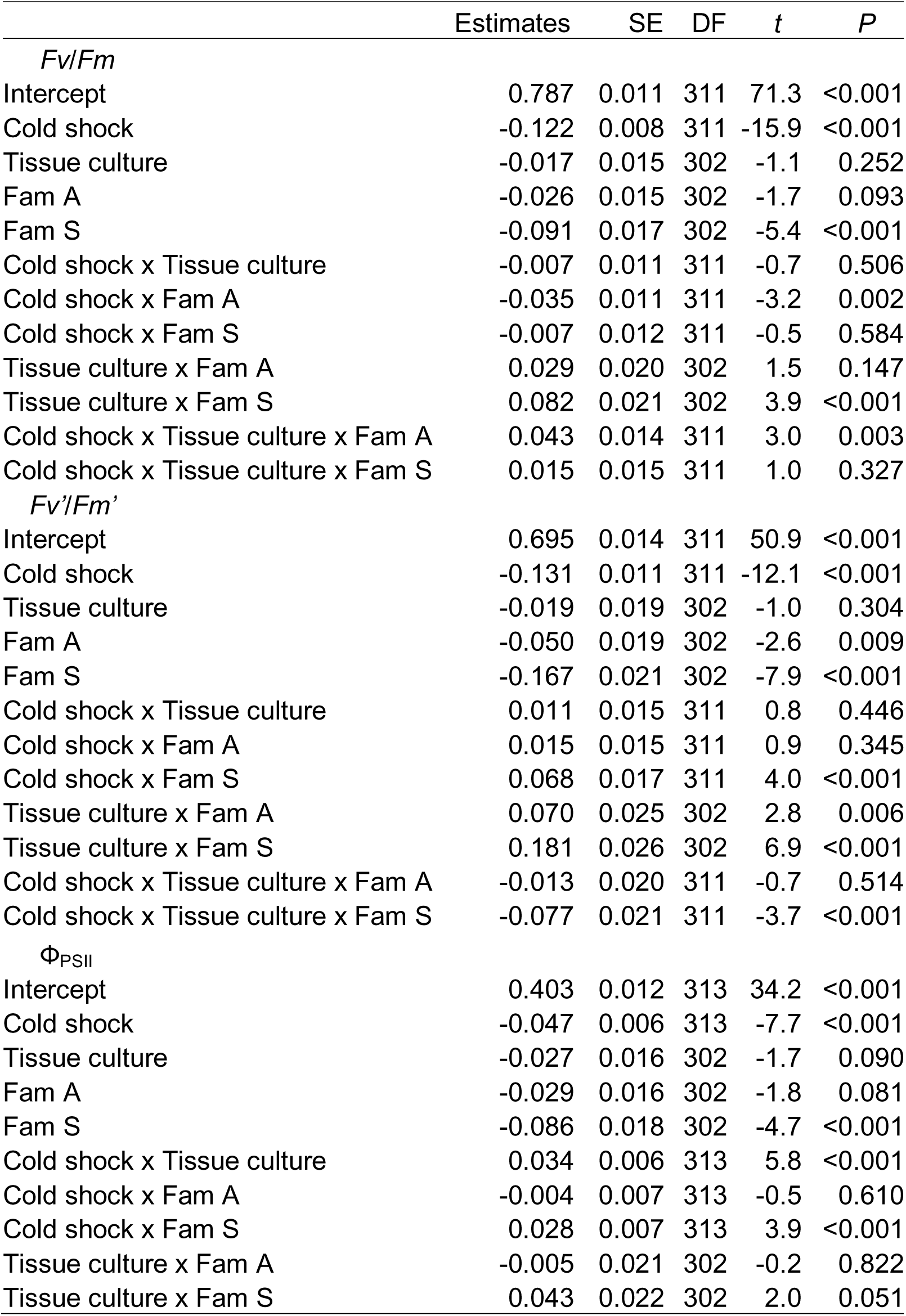
The best linear mixed models for *F*_*v*_/*F*_*m*_, *F*_*v*_’/*F*_*m*_’ and Φ_PSII_, based on AIC. Intercepts represent the mixture of background conditions, i.e. not cold shocked, not tissue cultured, and family Notsand. Fam A and Fam S refer to families Ardal and Sandfell, respectively.

### Effects of tissue culturing

We detected genotype–specific effects of tissue culture on *F*_*v*_/*F*_*m*_, *F*_*v*_*’*/*F*_*m*_*’* and Φ_PSII_ (Table 3). This is consistent with a previous report of a genotype–specific effect on callus characteristics (Glock 1989; Glock & Gregorius 1986). The three measured parameters of chlorophyll fluorescence (*F*_*v*_/*F*_*m*_, *F*_*v*_*’*/*F*_*m*_*’* and Φ_PSII_) all decreased after the cold treatment (Table 3), indicating a decrease in photosystem II activity, as reported in previous studies (Finazzi *et al*. 2006). A positive effect of interaction between tissue culture and cold shock for Φ_PSII_ suggests that tissue-cultured plants were less affected by cold shock than seedlings, and an interaction between tissue culture, cold shock and family suggests that the extent to which tissue-cultured plants were less affected by cold shock differed among families. Any differences among families in traits related to responses to the tissue-culture environment, including root-cutting, callus formation and growth on medium, might explain these observed interactions between tissue culture and family. This finding is consistent with the report that somaclonal variation is genotype–dependent and influenced by both the explant source and the tissue-culture protocol (George & Sherrington 1984), and a recent study that found that the effect of tissue culture on somatic mutations depended on genotype (Zhang *et al*. 2010). The effects of tissue culture-genotype interaction, however, were comparable to, or much smaller than the single effect of family (Table 3), indicating that such interactions would not mask the single effect of genotype. The interaction between tissue culture and family was much smaller in Φ_PSII_ (the range between maximum and minimum estimates was 0.043 – (–0.005) = 0.048, Table 2) than in *F*_*v*_/*F*_*m*_ (0.082 – 0 = 0.082) and *F*_*v*_*’*/*F*_*m*_*’* (0.181 – 0 = 0.181). The interaction between cold shock, tissue culture and family was detected only in *F*_*v*_/*F*_*m*_ and *F*_*v*_*’*/*F*_*m*_*’*. Also, the relative impact of among-genotype variance was smaller for Φ_PSII_ (8.5% of the total variance, Table 2) than *F*_*v*_/*F*_*m*_ (31.5 %) and *F*_*v*_*’*/*F*_*m*_*’* (10.9 %). These results imply that, although the maximum efficiencies of photosynthesis for dark‐ (*F*_*v*_/*F*_*m*_) and light-adapted leaves (*F*_*v*_*’*/*F*_*m*_*’*) were affected by tissue culturing in genotype– specific ways, the actual electron transport operating efficiency (Φ_PSII_) was less affected by tissue culture.

### Conclusion

Overall, we successfully detected among-genotype variance, with low somaclonal variation, indicating that the advantage of tissue culturing in generating genetically isogenic replicates exceeded its disadvantage in amplifying somaclonal variation in our study system. We detected interaction effects of tissue culture with genotype for an adaptively relevant trait, cold tolerance; however, such variation would not mask the single effect of genotype. Therefore, although one should carefully consider effects of tissue culturing when interpreting any results relying on the technique, tissue culturing is a useful method for obtaining genetically homogenous replicates in this, and probably other non-model organisms. It can provide critical additional power when studying phenotypes such as cold tolerance related to adaptation in natural environments, the variation in the phenotypes among families or populations, the reaction norms of a genotype or the QTLs accounting for phenotypes.

## Acknowledgements

We are grateful to Prof. M. Burrell for advice, Dr. P. Vergeer for providing seeds and Dr. C. Lilley and Ms. J. Hibbard for providing tissue culture protocols. This research was funded by the Natural Environment Research Council Post-Genomics and Proteomics programme (NE/C507837/1) in UK; the Special Coordination Funds for Promoting Science and Technology from the Ministry of Education, Culture, Sports, Science and Technology of the Japanese Government (MEXT); and research exchange program between Japan and UK by Japan Society for the Promotion of Science (10037611–000065).

## References

Alonso-Blanco C, Gomez-Mena C, Llorente F, et al. (2005) Genetic and molecular analyses of natural variation indicate CBF2 as a candidate gene for underlying a freezing tolerance quantitative trait locus in Arabidopsis. Plant Physiology 139, 1304–1312.

Baldi P, Pedron L, Hietala AM, La Porta N (2011) Cold tolerance in cypress (Cupressus sempervirens L.): a physiological and molecular study. Tree genetics & genomes 7, 79–90.

Bloomer RH, Lloyd AM, Symonds VV (2014) The genetic architecture of constitutive and induced trichome density in two new recombinant inbred line populations of Arabidopsis thaliana: phenotypic plasticity, epistasis, and bidirectional leaf damage response. BMC Plant Biology 14.

Bresson J, Vasseur F, Dauzat M, et al. (2015) Quantifying spatial heterogeneity of chlorophyll fluorescence during plant growth and in response to water stress. Plant Methods 11.

Charlesworth D, Mable BK, Schierup MH, Bartolome C, Awadalla P (2003) Diversity and linkage of genes in the self-incompatibility gene family in Arabidopsis lyrata. Genetics 164, 1519–1535.

Clauss MJ, Dietel S, Schubert G, Mitchell-Olds T (2006) Glucosinolate and trichome defenses in a natural Arabidopsis lyrata population. Journal of Chemical Ecology 32, 2351–2373.

Clauss MJ, Koch MA (2006) Poorly known relatives of Arabidopsis thaliana. Trends in Plant Science 11, 449–459.

Clauss MJ, Mitchell-Olds T (2006) Population genetic structure of Arabidopsis lyrata in Europe. Molecular Ecology 15, 2753–2766.

D'Amato F, Bayliss MW (1985) Cytogenetics of plant cell and tissue cultures and their regenerates. Critical Reviews in Plant Sciences 3, 73–112.

Davey MP, Burrell MM, Woodward FI, Quick WP (2008) Population-specific metabolic phenotypes of Arabidopsis lyrata ssp petraea. New Phytologist 177, 380–388.

Davey MP, Woodward FI, Quick WP (2009) Intraspecific variation in cold-temperature metabolic phenotypes of Arabidopsis lyrata ssp. petraea. Metabolomics 5, 138–149.

Dorn LA, Pyle EH, Schmitt J (2000) Plasticity to light cues and resources in Arabidopsis thaliana: Testing for adaptive value and costs. Evolution 54, 1982–1994.

Ehlert B, Hincha DK (2008) Chlorophyll fluorescence imaging accurately quantifies freezing damage and cold acclimation responses in Arabidopsis leaves. Plant Methods 4, 12.

El-Soda M, Malosetti M, Zwaan BJ, Koornneef M, Aarts MGM (2014) Genotype x environment interaction QTL mapping in plants: lessons from Arabidopsis. Trends in Plant Science 19, 390–398.

Finazzi G, Johnson GN, Dall'Osto L, et al. (2006) Nonphotochemical quenching of chlorophyll fluorescence in Chlamydomonas reinhardtii. Biochemistry 45, 1490–1498.

George EF, Sherrington PD (1984) Plant propagation by tissue culture Exegetics Ltd., Basingstoke.

Glock H (1989) Environmental effects on growth of tissue cultures of a woody Solanum species (Solanum laciniatum). Plant Science 62, 137–143.

Glock H, Gregorius HR (1986) Genotype–environment interaction in tissue cultures of birch. Theoretical and Applied Genetics 72, 477–482.

Gutteling EW, Riksen JAG, Bakker J, Kammenga JE (2007) Mapping phenotypic plasticity and genotype–environment interactions affecting life-history traits in Caenorhabditis elegans. Heredity 98, 28–37.

Heidel AJ, Clauss MJ, Kroymann J, Savolainen O, Mitchell-Olds T (2006) Natural variation in MAM within and between populations of Arabidopsis lyrata determines glucosinolate phenotype. Genetics 173, 1629–1636.

Heo J-Y, Feng D, Niu X, et al. (2014) Identification of quantitative trait loci and a candidate locus for freezing tolerance in controlled and outdoor environments in the overwintering crucifer Boechera stricta. Plant Cell and Environment 37, 2459–2469.

Hong S-W, Vierling E (2000) Mutants of Arabidopsis thaliana defective in the acquisition of tolerance to high temperature stress. Proceedings of the National Academy of Sciences of the United States of America 97, 4392–4397.

Jansen M, Gilmer F, Biskup B, et al. (2009) Simultaneous phenotyping of leaf growth and chlorophyll fluorescence via GROWSCREEN FLUORO allows detection of stress tolerance in Arabidopsis thaliana and other rosette plants. Functional Plant Biology 36, 902–914.

Joyce SM, Cassells AC, Jain SM (2003) Stress and aberrant phenotypes in in vitro culture. Plant Cell Tissue and Organ Culture 74, 103–121.

Khanal N, Moffatt BA, Gray GR (2015) Acquisition of freezing tolerance in Arabidopsis and two contrasting ecotypes of the extremophile Eutrema salsugineum (Thellungiella salsuginea). Journal of Plant Physiology 180, 3544.

Kuittinen H, Niittyvuopio A, Rinne P, Savolainen O (2008) Natural variation in Arabidopsis lyrata vernalization requirement conferred by a FRIGIDA indel polymorphism. Molecular Biology and Evolution 25, 319–329.

Kunin WE, Vergeer P, Kenta T, et al. (2009) Variation at range margins across multiple spatial scales: environmental temperature, population genetics and metabolomic phenotype. Proceedings of the Royal Society B-Biological Sciences 276, 1495–1506.

Kwon Y, Kim SH, Jung MS, et al. (2007) Arabidopsis hot2 encodes an endochitinase-like protein that is essential for tolerance to heat, salt and drought stresses. Plant Journal 49, 184–193.

Larkin PJ, Scowcroft WR (1981) Somaclonal variation—a novel source of variability from cell cultures for plant improvement. Theoretical and Applied Genetics 60, 197–214.

Lexer C, Welch ME, Durphy JL, Rieseberg LH (2003) Natural selection for salt tolerance quantitative trait loci (QTLs) in wild sunflower hybrids: Implications for the origin of Helianthus paradoxus, a diploid hybrid species. Molecular Ecology 12, 1225–1235.

Maxwell K, Johnson GN (2000) Chlorophyll fluorescence - a practical guide. Journal of Experimental Botany 51, 659–668.

McAusland L, Davey PA, Kanwal N, Baker NR, Lawson T (2013) A novel system for spatial and temporal imaging of intrinsic plant water use efficiency. Journal of Experimental Botany 64, 4993–5007.

Medeiros JS, Marshall DL, Maherali H, Pockman WT (2012) Variation in seedling freezing response is associated with climate in Larrea. Oecologia 169, 73–84.

Mishra A, Heyer AG, Mishra KB (2014) Chlorophyll fluorescence emission can screen cold tolerance of cold acclimated Arabidopsis thaliana accessions. Plant Methods 10.

Mitchell-Olds T (2001) Arabidopsis thaliana and its wild relatives: a model system for ecology and evolution. Trends in Ecology & Evolution 16, 693–700.

Murchie EH, Lawson T (2013) Chlorophyll fluorescence analysis: a guide to good practice and understanding some new applications. Journal of Experimental Botany 64, 3983–3998.

Quesada V, Garcia-Martinez S, Piqueras P, Ponce MR, Micol JL (2002) Genetic architecture of NaCl tolerance in Arabidopsis. Plant Physiology 130, 951–963.

Quick WP, Stitt M (1989) An examination of factors contributing to nonphotochemical quenching of chlorophyll fluorescence in barley leaves. Biochimica et Biophysica Acta 977, 287–296.

R Development Core Team (2008) R: A language and environment for statistical computing R Foundation for Statistical Computing., Vienna, Austria. http://www.R-project.org.

Sasaki E, Zhang P, Atwell S, Meng D, Nordborg M (2015) "Missing" G x E variation controls flowering time in Arabidopsis thaliana. Plos Genetics 11, e1005597.

Schierup MH, Bechsgaard JS, Christiansen FB (2008) Selection at work in selfincompatible Arabidopsis lyrata. II. Spatial distribution of S haplotypes in Iceland. Genetics 180, 1051–1059.

Shindo C, Bernasconi G, Hardtke CS (2007) Natural genetic variation in Arabidopsis: tools, traits and prospects for evolutionary ecology Annals Of Botany 99, 1043–1054.

Steponkus PL, Uemura M, Joseph RA, Gilmour SJ, Thomashow MF (1998) Mode of action of the COR15a gene on the freezing tolerance of Arabidopsis thaliana. Proceedings of the National Academy of Sciences of the United States of America 95, 14570–14575.

Stratton DA (1998) Reaction norm functions and QTL-environment interactions for flowering time in Arabidopsis thaliana. Heredity 81, 144–155.

Sultan S, Bazzaz F (1993) Phenotypic plasticity in Polygonum persicaria. I. Diversity and uniformity in genotypic norms of reaction to light. Evolution, 1009–1031.

Waitt D, Levin D (1993) Phenotypic integration and plastic correlations in Phlox drummondii (Polemoniaceae). American Journal of Botany, 1224–1233.

Woo NS, Badger MR, Pogson BJ (2008) A rapid, non-invasive procedure for quantitative assessment of drought survival using chlorophyll fluorescence. Plant Methods 4.

Wu RL (1998) The detection of plasticity genes in heterogeneous environments. Evolution 52, 967–977.

Xie HJ, Li H, Liu D, et al. (2015) ICE1 demethylation drives the range expansion of a plant invader through cold tolerance divergence. Molecular Ecology 24, 835–850.

Yap JS, Li Y, Das K, Li J, Wu R (2011) Functional mapping of reaction norms to multiple environmental signals through nonparametric covariance estimation. BMC Plant Biology 11.

Yuan F, Chen M, Leng BY, Wang BS (2013) An efficient autofluorescence method for screening Limonium bicolor mutants for abnormal salt gland density and salt secretion. South African Journal of Botany 88, 110–117.

Zhang JZ, Creelman RA, Zhu JK (2004) From laboratory to field. Using information from Arabidopsis to engineer salt, cold, and drought tolerance in crops. Plant Physiology 135, 615–621.

Zhang MS, Wang H, Dong ZY, et al. (2010) Tissue culture-induced variation at simple sequence repeats in sorghum (Sorghum bicolor L.) is genotype–dependent and associated with down-regulated expression of a mismatch repair gene, MLH3. Plant Cell Reports 29, 51–59.

Zhen Y, Ungerer MC (2008) Clinal variation in freezing tolerance among natural accessions of Arabidopsis thaliana. New Phytologist 177, 419–427.

